# Broiler Chickens and Early Life Programming: Microbiome transplant-induced cecal bacteriome dynamics and phenotypic effects

**DOI:** 10.1101/2020.08.13.240572

**Authors:** Gustavo A. Ramírez, Ella Richardson, Jory Clark, Jitendra Keshri, Yvonne Drechsler, Mark E. Berrang, Richard J. Meinersmann, Nelson A. Cox, Brian B. Oakley

## Abstract

The concept of successional trajectories describes how small differences in initial community composition can magnify through time and lead to significant differences in mature communities. For many animals, the types and sources of early-life exposures to microbes have been shown to have significant and long-lasting effects on the community structure and/or function of the microbiome. In modern commercial poultry production, chicks are reared as a single age cohort and do not directly encounter adult birds. This scenario is likely to initiate a trajectory of microbial community development that is significantly different than non-industrial settings where chicks are exposed to a much broader range of environmental and fecal inocula; however, the comparative effects of these two scenarios on microbiome development and function remain largely unknown. In this work, we performed serial transfers of cecal material through multiple generations of birds to first derive a stable source of inoculum. Subsequently, we compared microbiome development between chicks receiving this passaged cecal material, versus an environmental inoculum, to test the hypothesis that the first exposure of newly hatched chicks to microbes determines early GI microbiome structure and may have longer-lasting effects on bird health and development. Cecal microbiome dynamics and bird weights were tracked for a two-week period, with half of the birds in each treatment group exposed to a pathogen challenge at 7 days of age. We report that: i) a relatively stable community was derived after a single passage of transplanted cecal material, ii) this cecal inoculum significantly but ephemerally altered community structure relative to the environmental inoculum and PBS controls, and iii) either microbiome transplant administered at day-of-hatch appeared to have some protective effects against pathogen challenge relative to uninoculated controls. Differentially abundant taxa were identified across treatment types that may inform future studies aimed at identifying strains associated with beneficial phenotypes.

## Introduction

Poultry comprise an economically important global protein market and are common animal models used in basic and applied research. Since the middle of the last century, antimicrobial growth promoters (AGPs), in-feed antibiotics at sub-therapeutic concentrations, have been commonly used in commercial broiler chicken farming to improve feed conversion efficiency (Landers et al., 2012). Despite their proven efficacy, presumed to result from modulations of the gastrointestinal (GI) microbiome and host interactions (Danzeisen et al., 2011;Costa et al., 2017;Gadde et al., 2018), the specific mechanisms of action of AGPs remain largely unknown. In the last decade, concerns about antibiotic overuse and shifting consumer preferences have led to new regulatory guidelines and industry practices removing AGPs from feed in the E.U. and the U.S. Promising alternatives to AGPs include the modulation of the chicken GI microbiome with prebiotics such as starches in the diet, antimicrobials such as organic acids or phytochemicals, and mono- or mixed-culture probiotics, as reviewed elsewhere (Huyghebaert et al., 2011;Clavijo and Flórez, 2018;Yadav and Jha, 2019). While many of these alternatives to antibiotics have shown some efficacy compared to controls, re-capturing the performance benefits of AGPs remains an elusive goal. A better understanding of specific bacterial strains associated with desirable phenotypes could help identify effective probiotic alternatives to AGPs.

The establishment and population dynamics of the chicken GI microbiome have been fairly well-described (Lu et al., 2003;Oakley et al., 2014). Generally, immediately after hatching, colonization by environmental microbes and subsequent community succession results in hundreds of billions of bacterial cells per gram of GI content after just a few days (Ella and Barnes, 1979). Of particular interest within the GI tract are the ceca where the highest prokaryotic loads (Oakley et al., 2014) and the longest community residence times (Sergeant et al., 2014) are found. Importantly, the ceca are a major site for bacterial fermentations and the production of short-chain fatty acids [SCFAs; (Van der Wielen et al., 2000;Dunkley et al., 2007)]. SCFAs, including lactate, acetate, propionate, and butyrate, directly stimulate increases in absorptive surface area (Dibner and Richards, 2005), suppress the growth of zoonotic pathogens (Namkung et al., 2011), induce the expression of host-defense peptides (Sunkara et al., 2011), and modulate host epigenetic regulation (Canani et al., 2011).

One promising approach to better understand how specific GI bacterial taxa may influence growth performance and pathogen resistance in poultry is the use of microbiome transplants (MTs). Targeted modulation of the GI microbiome, particularly during early development, may significantly influence phenotypes in mature birds (Rubio, 2019). Work in mammalian models has shown that fecal transplants can affect host energy balance and weight gain (Turnbaugh et al., 2006;Ridaura et al., 2013). In chickens, transplantation of fecal excreta from healthy adult birds to newly hatched chicks has been shown to improve resistance against *Salmonella* (Siegerstetter et al., 2018). By inducing desirable phenotypes such as changes in body weight-gain or pathogen resistance via early life “metabolic programming” (Waterland and Garza, 1999), MTs can be used to infer which bacterial strains, consortia, or metabolic pathways may contribute to phenotypic effects observed in the host. MTs may affect host health via the competitive exclusion of potential pathogens, lowering community production of growth suppression metabolites, and/or improving host energy metabolism (Dibner and Richards, 2005;Yadav and Jha, 2019). Chickens in the natural environment are exposed to a wide diversity of microbes early in life from environmental sources and excreta from multi-age cohorts of birds. In contrast, chicks in typical commercial broiler production systems do not encounter adult birds and are reared as a single age cohort in relatively controlled conditions under modern biosecurity regimens. The importance of early-life exposures to microbes has been shown repeatedly for many host animals and humans; for example, cesarean-section vs. vaginal birth (Neu and Rushing, 2011), or breast-fed vs formula fed infants (Milani et al., 2017), but for chickens, how exposure to environmental versus host-derived microbial communities (e.g. FMT) shapes the microbiome, remains unknown.

Here, to better understand microbiome-host interactions and the effects of MTs, we assessed cecal microbiome dynamics of healthy broiler chicks, from hatching to 14 days post-hatch, administered one of three treatments: i) a community enriched from serial passages of cecal contents through multiple generations of chicks (CMT), ii) an environmental community obtained from commercial poultry litter (EMT), or iii) a phosphate buffer saline (PBS) control. At one week of age, approximately half of the chicks in each group were administered an oral gavage of a pathogen challenge. We report significant differential phenotypic effects elicited by specific MT treatments for weight gain and pathogen resistance. Further, we identify shifts in the cecal microbiome at the community- and strain-level and identify differentially abundant taxa across MT treatment types associated with observed phenotypes.

## Results

### 1. Community dynamics of serially passaged CMT

Community composition of the cecal microbiome transplants generally stabilized after a single passage (Figure 1A). Samples prior to the first serial passage were dominated (nearly 90% of all sequences) by the phylum Firmicutes, whereas, after one transfer, the phylum Bacteroidetes was dominant (Figure 1B). This shift in community composition at the phylum level after one transfer could be clearly seen in a stable Firmicutes to Bacteroidetes ratio after the first serial passage (Figure 1C). At the genus level, the community prior to the first serial passage was comprised primarily of Lactobacillus, Eubacterium, Faecalicoccus, and Anaerobacterium; whereas communities after one transfer were dominated by a few Bacteroides genera including Alistipes, Barnesiella, and Blautia (Figure 1D). Summaries of alpha diversity at the genus level showed significantly higher taxonomic richness prior to the first of the serial passages while all subsequent serial passages show lower and stable counts of observed genera (Figure 1E). Overall, despite some individual variability, frozen cecal material was significantly altered after the first passage and stable thereafter. This stable community derived from serial passages through young chicken ceca was subsequently used as the CMT inoculum in this study.

**Figure 1.**
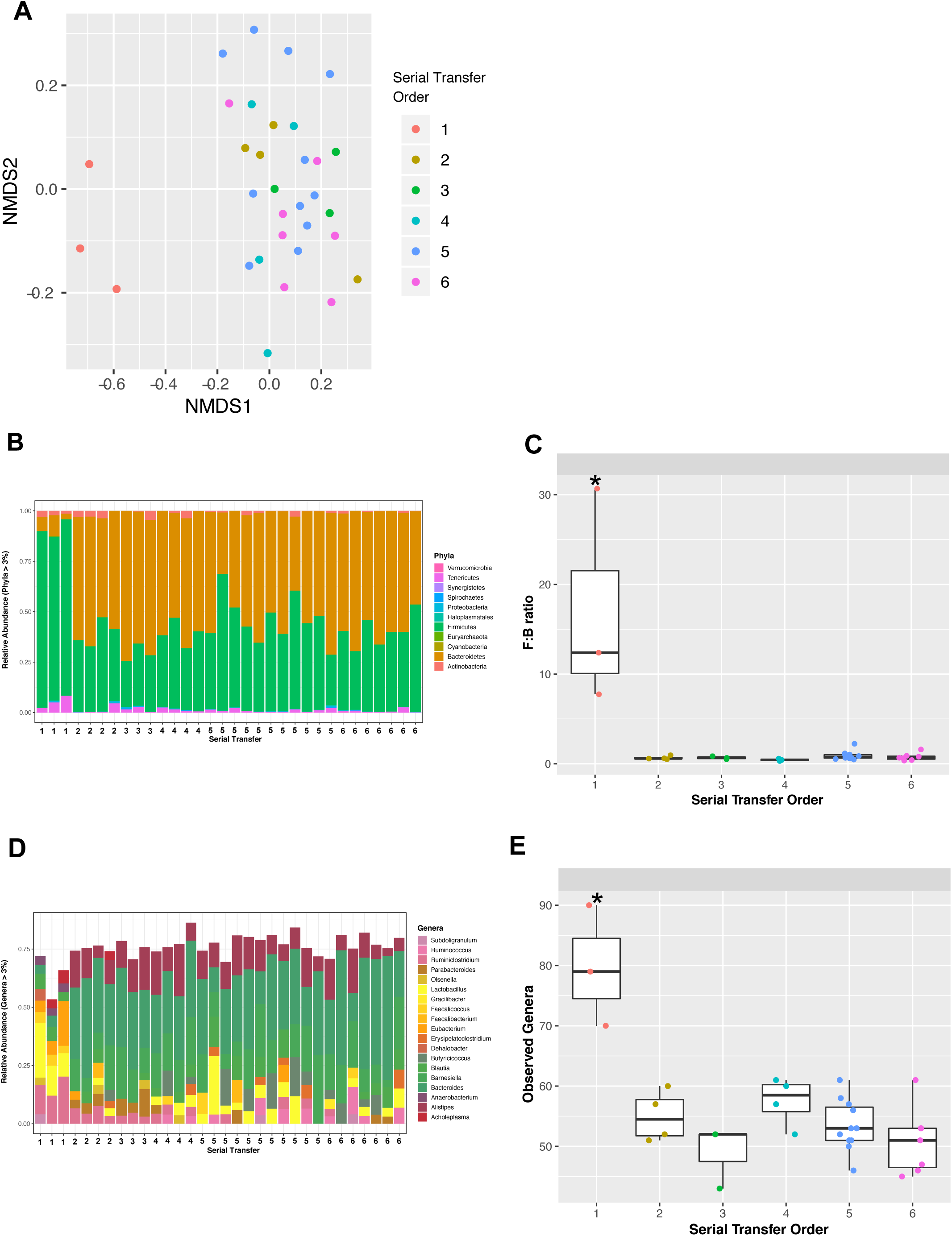
Microbiome analyses of serial transfer samples rarefied to even depth (n=850 sequences per sample). A) Ordination analysis color coded by serial transfer number. B) Phylum level community composition. C) Firmicutes:Bacteroidetes ratios for each serial passage. D) Genus level community composition. E) Number of observed genera as a function of serial transfer order (*: significantly different means, p < 0.05).

### 2. Bacterial community composition of gavage inocula

We used a simple factorial design to assess the effects of day-of-hatch microbiome transplant type (*i*.*e*.: EMT, CMT, and PBS) on cecal microbiome dynamics and pathogen resistance (Figure 2A). Community composition of the environmental and cecal-enrichment gavages (EMT and CMT treatments, respectively) differed drastically (Figure 2B). Over 98% of the sequences recovered from the EMT gavage belong to the phylum Firmicutes, primarily the genus *Lactobacillus*. In contrast, at the phylum-level, the CMT gavage community was predominantly (>75%) comprised of the phylum Bacteroidetes with the remainder (< 25%) of sequences classified as Firmicutes. At the genus-level, the CMT gavage was more diverse than the EMT gavage with the Bacteroidetes genera *Alistipes, Bacteroides*, and *Barnesiella* representing approximately 75% of the CMT community (Figure 2B).

**Figure 2.**
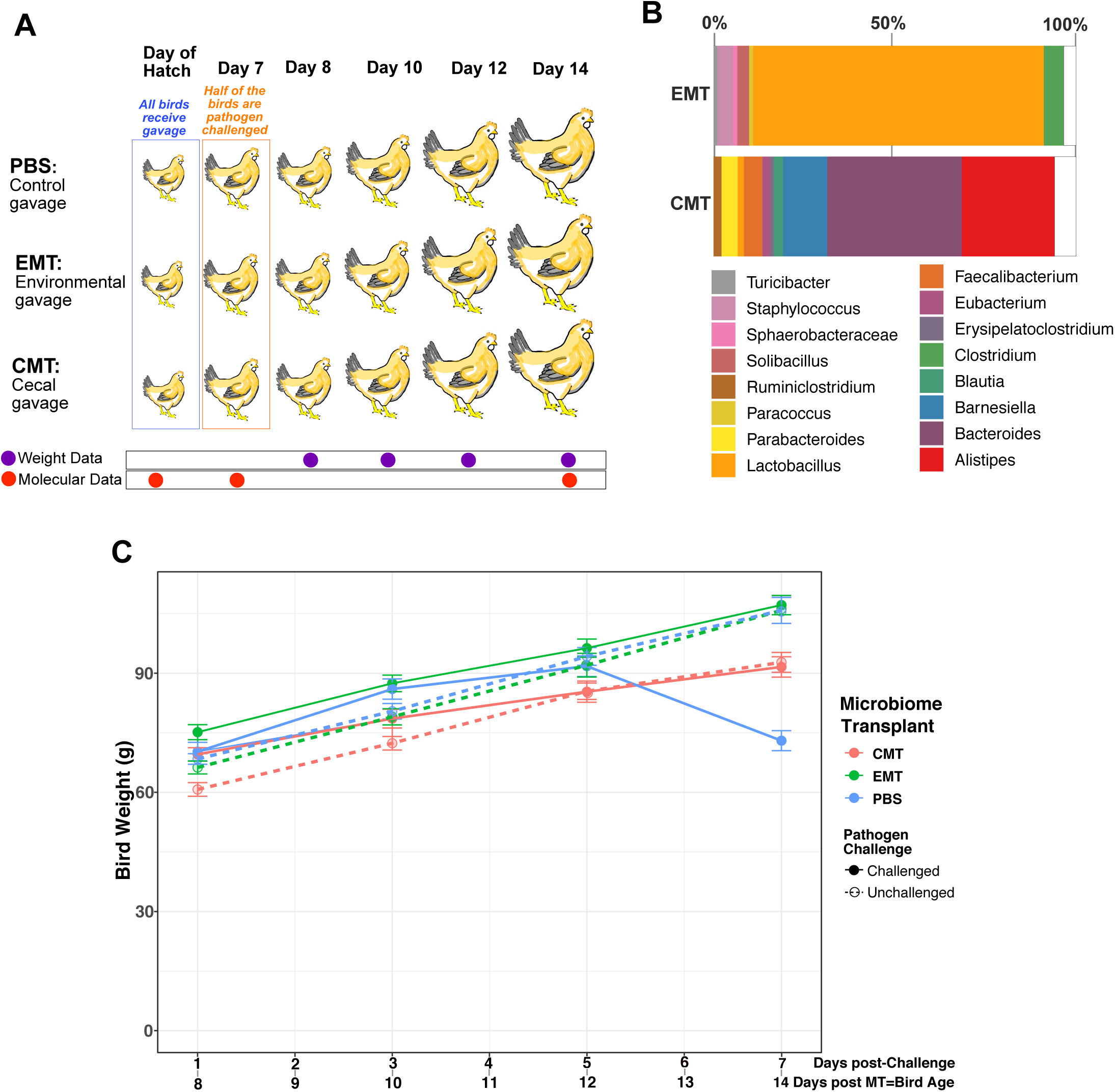
A) Schematic of the pooled cross-sectional study design for assessing the combined influence MT type (PBS, EMT, CMT) and pathogen challenge status (challenged vs. not-challenged). MT (via oral gavage) and pathogen challenge administration, both experimental variables, are time-stamped and depicted in blue and orange fonts, respectively. Longitudinal cross-sectional data collection for cecal molecular analyses and panel data collection for bird weight time series are depicted by red and orange purple, respectively. B) Bacterial community composition at the genus-level for gavages used to administer EMT and CMT in day-of-hatch chicks. C) Time series results for bird weight as a function of MT type and pathogen challenge status for birds age 8 through 14 days.

### 3. Bird Weight as a function of treatment group and pathogen challenge

Body weight differences across treatment groups were only significantly different at d14 post-hatching (Figure 2C, Table 1). In the non-challenged group at d14, weight distributions significantly differed as a function of the type of day-of-hatch MT received; EMT and PBS recipients were significantly heavier relative to birds that received a CMT. Interestingly, in the pathogen-challenged group at d14, significant differences were observed as a function of receiving either CMT or EMT at day-of-hatch relative to PBS controls. The PBS gavage (negative MT control) recipients lost approximately 20% of their average body weight between 12 and 14 days of age (5-7 days post-challenge) and at day 14 of age were significantly lighter than MT (EMT and CMT) recipients. Also, at d14 of age, EMT recipients were significantly heavier than CMT recipients.

**Table 1.** Weight data replicates used to produce Figure 2C. The same 53 birds had their weight in two-day intervals at the following post-microbiome transplant (bird age) dates and data was tabulated as a function of gavage type and pathogen challenge status (NC for not challenged and C for challenged).

|  | **CMT** | **EMT** | **PBS** |
| --- | --- | --- | --- |
| **Day 8** | n= 20 (11NC, 9C) | n= 15 (8NC, 7C) | n= 18 (10NC, 8C) |
| **Day 10** | n= 20 (11NC, 9C) | n= 15(8NC, 7C) | n= 18(10NC, 8C) |
| **Day 12** | n= 20 (11NC, 9C) | n= 15(8NC, 7C) | n= 18(10NC, 8C) |
| **Day 14** | n= 20 (11NC, 9C) | n= 15(8NC, 7C) | n= 18(10NC, 8C) |

### 4. Alpha-diversity

The number of observed taxa (genus- and 99% OTU-level) was lowest in 1-day old birds for all treatment groups (Figure 3, A & B). However, significantly more taxa at the genus and 99% OTU levels were observed at d1 for birds administered a CMT relative to the EMT treatment or PBS controls (Figure 3, A & B). From day 1 to day 7, significant increases in observed taxa occurred for all treatment groups (Figure 3). Subsequently, for birds that did not undergo a pathogen challenge, there were no significant differences in genus- or OTU-level richness between bird age 7 and 14 days (Figure 3, A & B). For birds that were pathogen challenged at 7 days of age, a significant decrease in OTU-level richness at 14d relative to 7d was observed in the group that received a day-of-hatch CMT (Figure 3D). A day-of-hatch CMT administration generally resulted in higher OTU richness at d7 versus d14 for both the non-challenged and pathogen-challenged groups; however, these observations were only statistically significant in the challenged group (Figure 3, B & D).

**Figure 3.**
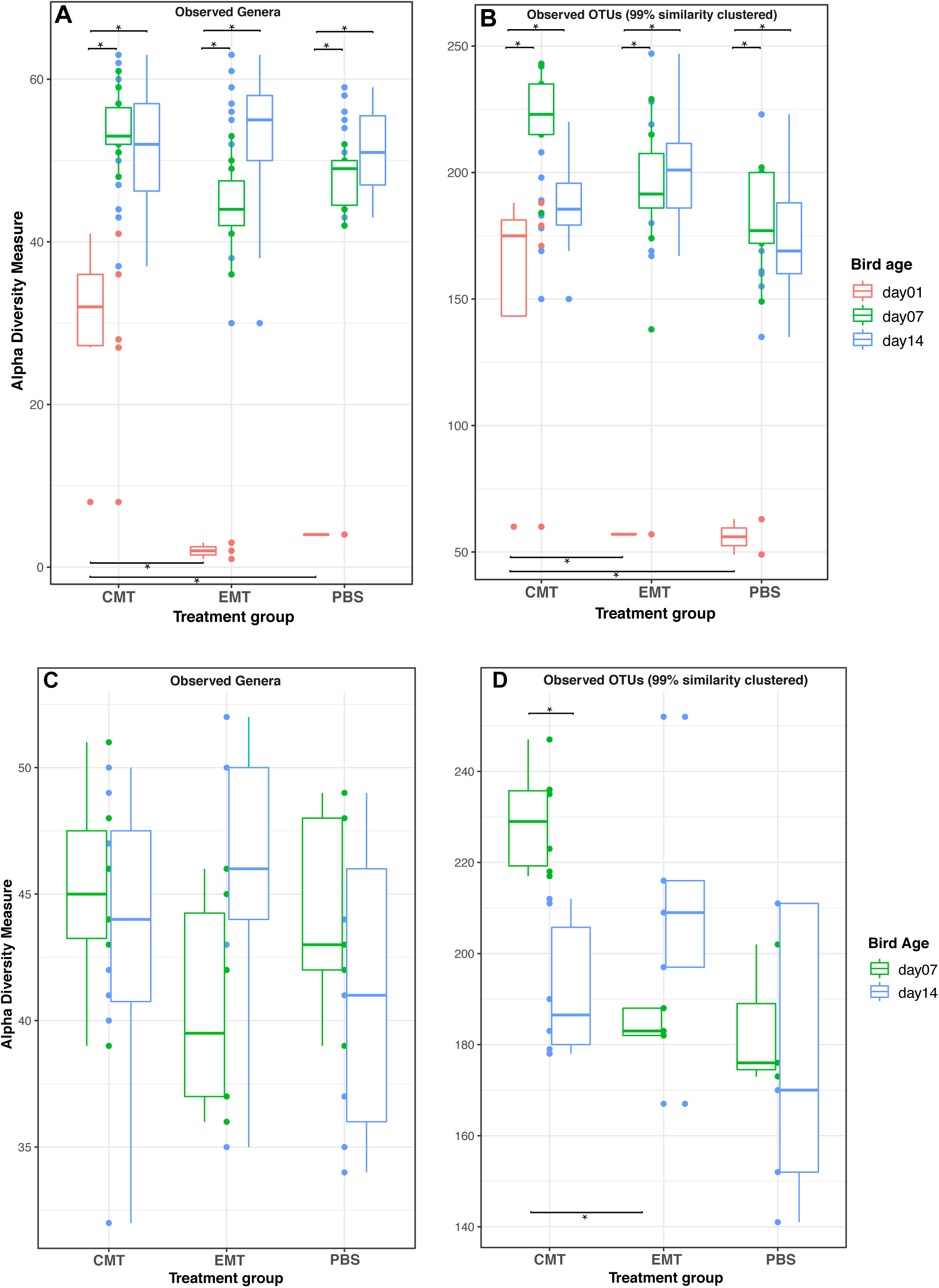
Community richness summary for each experiment group as a function of time (bird age in days). Only taxa with abundances greater than 5 in the dataset and samples with 1000 sequences are retained. All samples were rarefied to even depth A) Operational taxonomic units defined at the Genus-level (n= 1012 per sample) for non-pathogen challenged group. B) Operational taxonomic units defined at the 99% sequence similarity-level (n=1044 per sample) for non-pathogen challenged group. C) Operational taxonomic units defined at the Genus-level (n= 1012 per sample) for pathogen challenged group. D) Operational taxonomic units defined at the 99% sequence similarity-level (n=1044 per sample) for pathogen challenged group. Horizontal bars with asterisks denote significant differences between comparison pairs (student t-test, alpha = 0.05). Significant differences within MT groups and between MT groups are depicted at the top and bottom of the figure, respectively.

### 5. Beta-diversity

Cecal communities of 1-day old birds (1d) showed few distinct patterns but CMT recipients generally clustered close to the CMT gavage itself along positive axis 1 and 2 values (Figure 4A). Cecal communities from EMT and PBS recipients and the EM gavage spread along the range of axis 2 but were largely confined to negative axis 1 values (Figure 4A). By 7 days of age (d7), cecal communities from birds that received a PBS gavage instead of a microbiome transplant were most similar to each other and generally clustered along negative axis 1 values (Figure 4, B & D). Cecal communities of CMT or EMT recipients also clustered together and were more similar to the CMT than the EMT gavage community (Figure 4, B & D). By 14 days of age (d14), community distinctions among treatments collapsed and no discernable patterns associated with MT type were observed (Figure 4, C & E).

**Figure 4.**
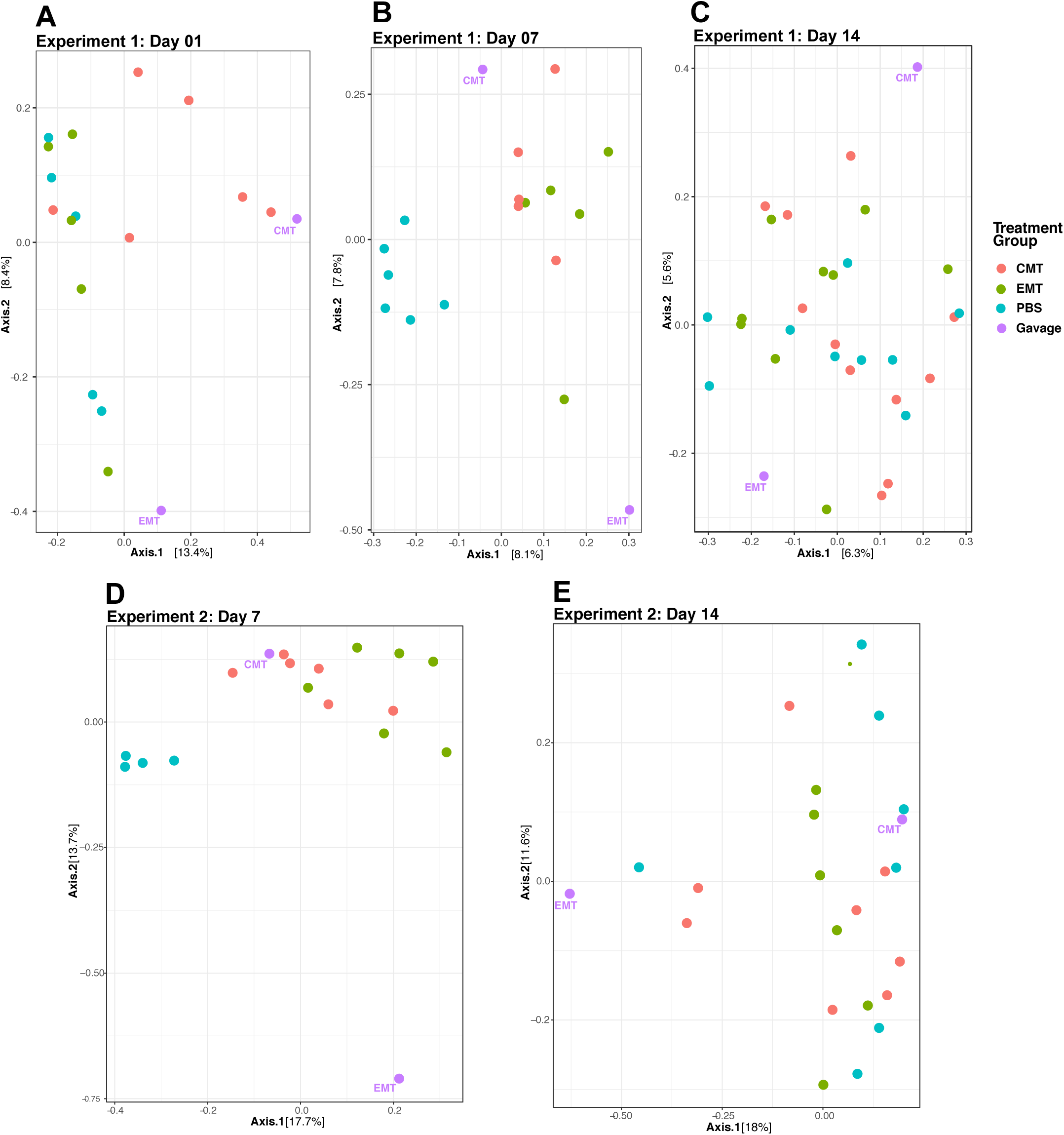
A-C: Ordinations plots depicting community composition for unchallenged bird group of each treatment group as a function of time (bird age in days). D & E: Ordinations plots depicting community composition for challenged birds of each treatment group as a function of time (bird age in days).

### 6. Differentially abundant taxa in MTs relative to PBS controls in 7-day old chicks

#### 6.1.1 Unchallenged Birds: EMT

A total of 9 OTU lineages, belonging to three genera within the phylum Bacteroidetes, exhibited significant differences in abundance in cecal communities from unchallenged birds that received EMTs compared to PBS controls (Figure 5A). These OTUs were classified as members of the *Barnesiella, Parabacteroides*, and *Alistipes* genera (Figure 5A).

**Figure 5.**
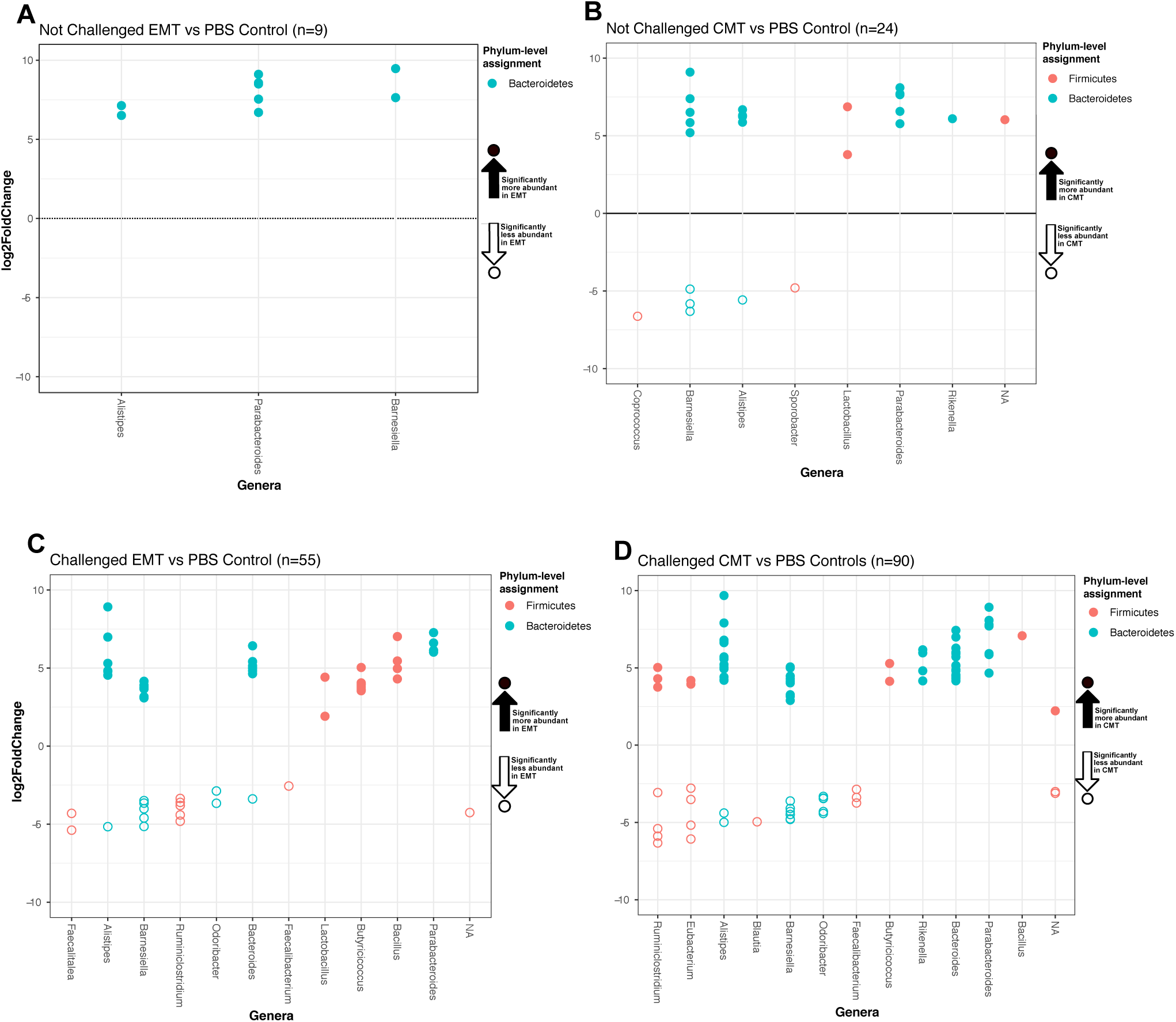
Taxa exhibiting significant differences in abundance following MT treatments relative to PBS controls in the cecal communities of 7-day old birds. The x-axis shows taxonomic assignment at the genus-level for individual OTU depicted as circles. Circle color depicts phylum-level taxonomic assignments. The y-axis shows the differential Log2-fold abundance change for each taxon. Open circles represent OTUs that are significantly (Wald Test, alpha=0.05) less abundant in MT data relative to PBS. Closed circles represent OTUs that are significantly (Wald Test, alpha=0.05) more abundant in MT data relative to PBS. See Supplemental Materials for a comprehensive list of differentially abundant OTU IDs and fasta sequences. A) Not challenged group: Significant differences in EMT relative to controls. B) Not challenged group: Significant differences in CMT relative to controls. C) Pathogen challenged group: Significant differences in EMT relative to controls. D) Pathogen challenged group: Significant differences in CMT relative to controls.

#### 6.1.2 Unchallenged Birds: CMT

A total of 24 OTU lineages, belonging to either the Firmicutes or Bacteroidetes exclusively, were significantly differentially abundant in cecal communities from unchallenged birds that received CMT relative to PBS controls (Figure 5B). Specifically, 18 OTUs were significantly more abundant in CMT versus PBS treatments (Figure 4B). These OTUs were classified within the following genera: *Rikenella, Parabacteroides, Lactobacillus, Alistipes*, and *Barnesiella* (Figure 5B). Five OTUs classified as *Coprococcus, Barnesiella, Alistipes* and *Sporobacter* were significantly less abundant in CMT versus PBS treatments (Figure 5B). Interestingly, two genera, *Alistipes* and *Barnesiella*, had OTUs that were both significantly more and less abundant in cecal communities of CMT recipients relative to PBS controls (Figure 5B).

#### 6.2.1 Pathogen Challenged Birds: EMT

A total of 54 OTU lineages, belonging to either the Firmicutes or Bacteroidetes, exhibited significant differences in abundance in cecal communities from pathogen challenged birds in the EMT group versus PBS controls (Figure 5C). Specifically, thirty-six and nineteen OTU lineages were significantly more and less abundant, respectively, in cecal communities from EMT recipients relative to PBS controls. All OTUs classified as *Lactobacillus, Butyricicoccus, Bacillus*, and *Parabacteroides*, were significantly enriched in EMT relative to PBS controls. All OTUs classified as *Faecalitalea, Barnesiella, Odoribacter*, and *Faecalibacterium* were significantly less abundant in cecal communities from birds that received an EMT relative to PBS controls. Interestingly, three genera (*Alistipes, Barnesiella*, and *Bacteroides*) contained some OTUs that were significantly enriched and some that were significantly less abundant in cecal communities of EMT recipients relative to controls (Figure 5C).

#### 6.2.2 Pathogen Challenged Birds: CMT

A total of 90 OTU lineages, belonging to either the Firmicutes or Bacteroidetes, exhibited significant differences in abundance in cecal communities from pathogen challenged birds that received a CMT compared to PBS controls (Figure 5D). 61 and 29 OTU lineages were significantly more abundant or less abundant, respectively, in cecal communities from CMT recipients relative to PBS controls. All OTUs classified as *Butyricicoccus, Rikenella, Bacteroides, Parabacteroides*, and *Bacillus*, were significantly enriched in CMT relative to PBS controls. All OTUs classified as *Odoribacter, Blautia*, and *Faecalibacterium*, were significantly less abundant in CMT relative to PBS controls. Four genera (*Alistipes, Barnesiella, Ruminiclostridium*, and *Eubacterium*) contained some OTUs that were significantly enriched and some that were significantly less abundant in cecal communities of CMT recipients relative to PBS controls (Figure 5D).

### 6.3 Taxa Differentially Abundant in Both Challenged and Unchallenged Groups

A total of 178 OTU lineages exhibited significant differences in relative abundance between birds that received a MT (EMT or CMT) versus PBS controls (Figure 6A). 125 and 13 of these OTUs were observed exclusively in challenged and unchallenged groups, respectively. Twenty differentially abundant OTUs, all classified as Bacteroidetes, were observed in both pathogen-challenged and unchallenged groups. Interestingly, these 20 OTUs exhibit similar trends in magnitude and fold change direction as a function of MT administration in both pathogen-challenged and unchallenged groups even though these were independent experimental cohorts (Figure 6B).

**Figure 6.**
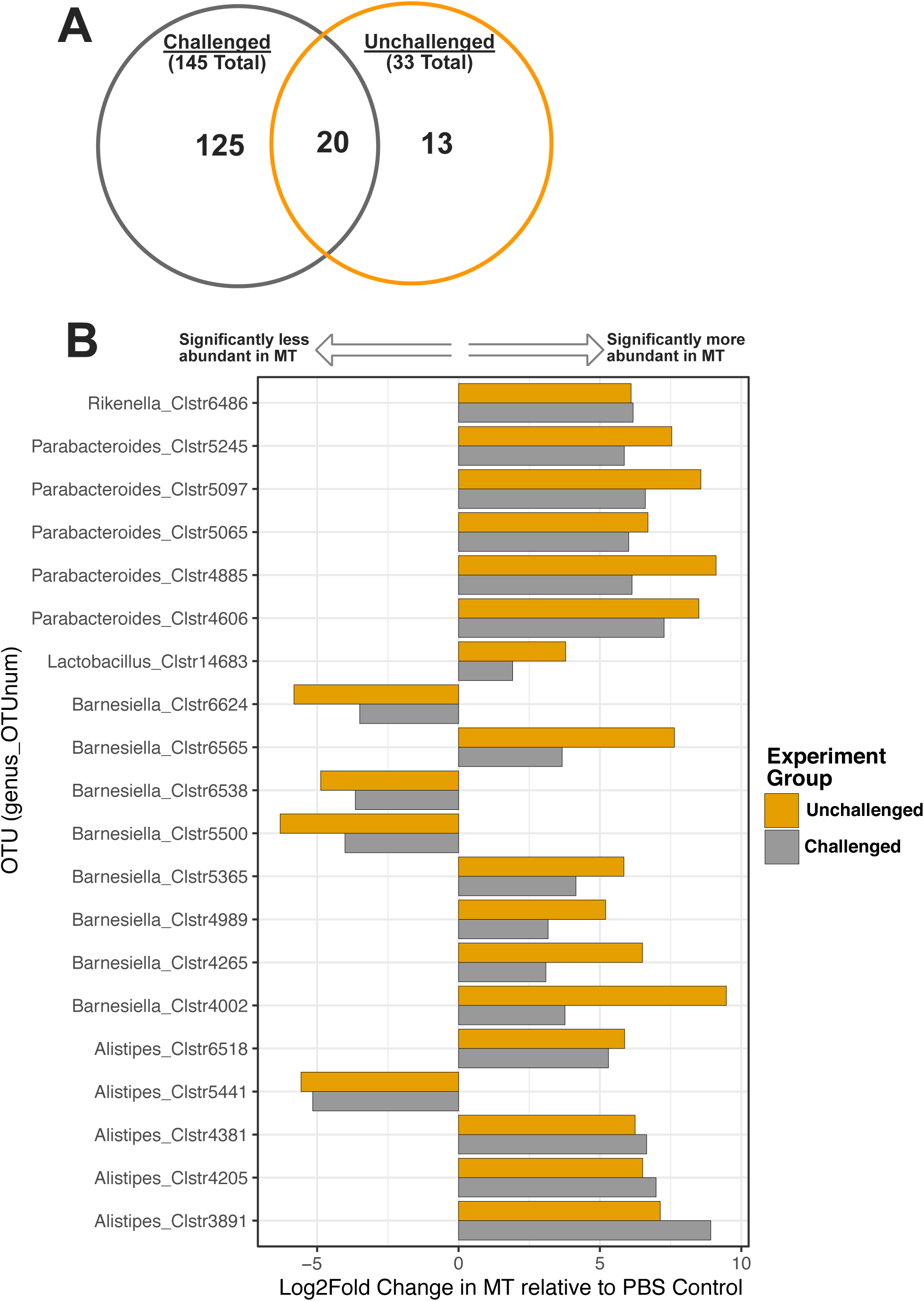
Observed abundances of differentially abundant taxa present in both pathogen-challenged and unchallenged experiment groups. The x-axis shows the differential Log2-fold abundance change for each OTU observed per experiment group (challenge and unchallenged abundances summarized in gray and orange, respectively). The y-axis shows taxonomic assignment at the genus-level for each OTU.

## Discussion

Applying the conceptual framework of successional trajectories (Fastie, 1995), similar to the concept of “early life programming” (Rubio, 2019), we hypothesized that first exposure of newly hatched chicks to environmental microbes determines early GI microbiome structure and may have long-lasting effects on bird health and development. To test this hypothesis, we tracked cecal microbiome dynamics and pathogen resistance of broiler chicks that received complex microbiome transplants at day-of-hatch. To compare the effects of very different first microbial exposure starting points, we compared a stable inoculum derived from serial passages of cecal material to a complex environmental community derived from used poultry litter and sterile PBS controls. To assess if early microbial exposure influences resistance to pathogenic infection, we performed this study on two bird panels, one that was pathogen challenged at 7d of age and one that was not pathogen challenged (Figure 2A).

### Microbiome dynamics through serial passages of cecal material

To obtain a transplant community inoculum selected by the cecal environment of broiler chicks, we serially transplanted cecal material from 14-day-old birds to newly hatched chicks. When the chicks reached 2 weeks of age, cecal contents were harvested and transplanted to a new batch of chicks. This serial passaging was repeated for five generations of chicks. We hypothesized that environmental filtering (Szekely and Langenheder, 2014) would result in an overall reduction in community richness with each serial transfer of cecal material and eventually lead to a stable microbial cohort consistently sorted by environmental and host-mediated factors. After just one passage, a relatively stable inoculum was derived (Figure 1). After the first serial passage, the starting inoculum had changed significantly in community diversity and composition from Firmicutes to Bacteroidetes dominance and remained relatively stable thereafter (Figure 1 C-E). These results suggest that a taxonomic subset of a community is quickly selected in a deterministic fashion by the host. We speculate that, given its relative stability, the selected community should be beneficial to the host.

### Either CMTs or EMTs enhance resistance to pathogen infection

We observed two significant effects of day-of-hatch MT on bird weight. First, in unchallenged birds, day-of-hatch EMT administration had no effect on weight while CMT administration led to significantly lower bird weight relative to controls (Figure 2C). In pathogen challenged birds, administration of either MT type resulted in higher bird weight relative to controls; however, birds administered the EMT gavage were significantly heavier than CMT recipients (Figure 2C). These observations lend credence to the notion that MT-elicited modulations of the GI flora, are both a consequence of host genetics and health status (Schokker et al., 2015), and also a cause of changes in host phenotype. Because EMT rather than CMT administration resulted in increased weight gain, independent of pathogen challenge status, we concluded that gavage composition drives phenotypic outcomes and that EMT inoculation alone may be sufficient to produce desirable phenotypes. The EMT gavage was largely comprised of Firmicutes lineages assigned as *Lactobacillus* spp. while the CMT was primarily comprised of Bacteroidetes lineages within the *Alistipes, Bacteroides*, and *Barnesiella* genera. Notably, despite being sourced from used commercial poultry litter, the EMT composition (predominantly Firmicutes, Figure 2B) differs from previously reported communities of chicken feces [predominantly Proteobacteria (Siegerstetter et al., 2018)]. Generally, a high prevalence of Firmicutes in the broiler GI tract is associated with beneficial immunomodulation (LeBlanc et al., 2013;Oakley and Kogut, 2016). *Lactobacillus* spp. are common probiotics that have been shown to enhance energy metabolism (LeBlanc et al., 2013), and inhibit colonization of *Campylobacter jejuni* in broilers (Neal-McKinney et al., 2012). Together, these factors may explain our observations that EMT treatment consistently resulted in higher bird weight relative to CMT. However, we note that the CMT gavage, comprised primarily of Bacteroidetes lineages, also resulted in increased weight gain relative to controls in pathogen challenged birds. This suggests that Firmicutes dominance (*Lactobacillus* spp., specifically) is not the sole determinant of the phenotypic effects elicited by both MT types in pathogen challenged birds. Overall, enhanced resistance to pathogen infection, inferred from weight gain, during early development (< 2 weeks of age) appears to be a global benefit conferred by administration of day-of-hatch MT (EMT and CMT) in broilers.

### MT-induced bacteriome dynamics

Early life microbiome status plays a critical role in establishing immune functions in murine (Cahenzli et al., 2013) and chicken models (Schokker et al., 2015). We report rapid increases in community richness between 1d and 7d independent of MT type administered at day-of-hatch and pathogen-challenge status, however, richness generally remained stable between 7d and 14d. This corroborates previous work suggesting the rapid (within less than a week post-hatching) establishment of taxonomically rich GI communities (Apajalahti et al., 2004). Interestingly, pathogen-challenged birds at 7d had significantly more diverse cecal communities if a CMT gavage was administered at day-of-hatch (Figure 3D), however, no additional effects of either MT treatment on bacterial community richness were observed. Enrichment of *Lactobacillus* spp. and a concurrent drop in alpha-diversity have been reported in chicken ceca of birds receiving Virginamycin as a prophylactic AGP (Costa et al. 2017); here, MT administration generally led to higher observed community richness relative to controls, however, these observations were not statistically significant (Figure 3). Ordination analyses of 7d cecal communities show compositional differences between birds that received MTs relative to controls in both pathogen-challenged and non-challenged birds (Figure 4 B & D). Given that differences in bird weight as a function of administered MT were observed at 14d, the microbial community clustering at 7d, where both CMT and EMT communities are similar to each other and dissimilar to controls, is particularly intriguing. Both MT types altered the cecal microbiome relative to controls prior to the observed phenotypic differences. These short-lived patterns in cecal bacteriome structure completely dissipate by 14d (Figure 4E) but may have had longer lasting effects on bird phenotype since both CMT and EMT recipients exhibit weight trajectories that were unaffected by pathogen challenge (Figure 2C). Overall, we show that ephemeral GI microbial community states specifically elicited by MTs administration early in a bird’s life may result in longer-lasting phenotypes. The mechanisms underlying this observation may involve immunological programming (Schokker et al., 2015;Oakley and Kogut, 2016) and are worthy of further investigations.

### Differentially abundant lineages

To better understand the potential mechanisms of action of MTs, we identified taxa that were significantly differentially abundant between MTs and control communities at 7d (Figure 5). In non-pathogen challenged birds, significantly higher abundances of 9 lineages belonging to the *Barnesiella, Parabacteroides*, and *Alistipes* genera were observed in the EMT treatments relative to controls at 7d (Figure 5A). The differential abundance of these taxa at 7d did not result in significant differences in bird weight at 14d (Figure 2C). Conversely, day-of-hatch CMT administration did result in lower bird weights at 14d relative to controls (Figure 2C), and thus taxa that differed significantly between the CMT and control communities at 7d (Figure 5A), may represent specific lineages implicated in longer term phenotypic outcomes. At 7d, taxa significantly less abundant in CMT communities relative to controls were *Coprococcus, Barnesiella, Alistipes*, and *Sporobacter* spp. while *Lactobacillus, Parabacteroides*, and *Rikenella* spp. OTUs were significantly more abundant relative to controls (Figure 5B). Other studies have reported *Coprococcus* spp., a butyrate-producing genera (Pryde et al., 2002), enriched in chicken ceca in response to AGP treatment (Danzeisen et al., 2011). A depletion of Coprococcus at 7d in the CMT treatment may lead to lower production of SCFAs which are well-described as key microbially-produced metabolites mediating host GI tract health, resulting in lower bird weight by 14d in our study. *Lactobacillus* spp. have been implicated in improved feed conversion ratios (Torok et al., 2011) and reduced mortality (Timmerman et al., 2006) in broilers and are thus generally considered beneficial probiotics (Bai et al., 2013). Despite the relative enrichment of *Lactobacillus* spp., birds in the CMT group ultimately experienced less weight gain relative to controls. Remarkably, the 9 lineages that were significantly more abundant in the 7d cecal communities of EMT recipients were also significantly more abundant in CMT recipient communities, even though the EMT and CMT treatments were derived and administered independently. These taxa may represent a core transplant microbiome, perhaps part of a consortium. Based on performance outcomes, the differentially abundant lineages in the CMT comparison, a total of 18 OTUs, should be considered potential performance-related phylotypes. In contrast, the subset of 9 lineages differentially abundant in the EMT comparison were not associated with any significant phenotypic differences. Together, these observations highlight specific OTU lineages that are differentially abundant across MTs and controls at critical points in early cecal community establishment and may provide clues to disentangle the complex links between broiler microbiome modulation and desirable phenotypes.

In pathogen challenged birds, day-of-hatch administration of a CMT or EMT gavage resulted in significantly higher bird weight relative to controls at 14d (Figure 2C). Taxa that were differentially abundant in both the CMT and EMT treatments at 7d compared to controls include: i) increases in OTUs assigned to the *Bacillus, Parabacteroides*, and *Butyricicoccus* genera, ii) depletion of OTUs assigned to the *Odoribacter*, and *Faecalibacterium* genera, iii) and increases and decreases in OTUs within the genera *Barnesiella* and *Alistipes* (Figure 5 C &D). Both *Bacillus* and *Butyricicoccus* ssp. are currently used as probiotics that have been shown to reduce heat stress-associated inflammatory responses (Wang et al., 2018) and confer protection against necrotic enteritis (Eeckhaut et al., 2016), respectively, in broiler chickens. Interestingly, despite being a common lineage recovered from chicken feces, here *Parabacteroides* spp. is significantly enriched along with *Bacillus* and *Butyricicoccus* spp., suggesting its potential as a possible probiotic. *Faecalibacterium* spp. have been repeatedly associated with positive health outcomes in humans (Sokol et al., 2008;Miquel et al., 2013) and have also been inversely correlated with expression of pro-inflammatory cytokines in broiler chickens (Oakley and Kogut, 2016). *Odoribacter* spp. decreases in cecal communities have been associated with butyric acid supplementation in chicken diets (Bortoluzzi et al., 2017). Together, these observations suggest that increases in abundance and/or activity of butyrate-producing taxa, such as *Faecalibacterium* and *Butyricicoccus* spp., may in fact dictate community dynamics and host-microbiome activities by generating fermentative metabolites and perhaps influence phenotypes later in life. Interestingly, we observed multiple genera (*Alistipes, Barnesiella, Bacteroides, Ruminiclostridium*, and *Eubacterium*) with OTUs that were both positively and negatively associated with experimental treatment and phenotype, reinforcing existing dogma that ‘strains matter’, i.e. specific bacterial strains can elicit significantly different phenotypes (). We note that in pathogen challenged birds, day-of-hatch MT administration yielded significantly higher bird weights relative to controls, however, the highest weight gains were observed in EMT recipients (Figure 1C). Two OTU lineages of *Lactobacillus* spp. were significantly more abundant in the EMT recipients at 7d relative to controls. Butyrate producers are known to cross-feed with lactic acid produced by *Lactobacillus* spp. (De Maesschalck et al., 2015) and the significant co-enrichment of *Lactobacillus* and, for example, *Butyricicoccus* spp. in the 7d cecal community of EMT recipients relative to controls, not observed in CMT recipients, suggests that the observed benefits of MT administration may result from enhanced cecal SCFA production.

### Conclusions

To advance our knowledge of microbiome-induced modulation of host health outcomes, microbiome transplants can provide predictive and testable guidance by identifying specific taxa that are differentially represented between treatments. Here we used MTs to better understand microbiome establishment from diverse inocula and to identify specific strains associated with pathogen resistance. Our results show that i) a relatively stable community was derived after a single passage of transplanted cecal material, ii) this cecal inoculum significantly but ephemerally altered community structure relative to the environmental inoculum and PBS controls, and iii) either microbiome transplant administered at day-of-hatch appeared to have some protective effects against pathogen challenge relative to uninoculated controls. We identify lineages that significantly differ in abundance in cecal contents from birds treated with MTs at day-of-hatch relative to controls that may drive observed phenotypic effects. These results suggest that environmental exposure to used poultry litter may provide an effective inoculum that could protect against pathogens and identifies specific taxa that may be responsible for this effect.

## Materials and Methods

### Microbiome Transplant Source Materials

The CMT source material was developed as follows: Frozen cecal material from 6 week-old broiler chickens was reconstituted by diluting 3:1 (w:v) in PBS and 0.2 mL administered via oral gavage to ten day-of-hatch chicks. When these chicks reached 2 weeks of age, their cecal contents were similarly prepared and administered to the next set of ten chicks. This serial passaging was repeated for five sets of chicks, with 10 chicks belonging to each group for a total of 50 birds. Chicks in each cohort were housed together. Cecal contents from each bird were sequenced as described below. The cecal contents from the final 10 birds were suspended 3:1 in PBS, pooled, and used immediately as the CMT inoculum. The EMT source material was generated from built up litter collected from a commercial poultry operation mixed 3:1 (w:v) in PBS and also provided as an oral gavage of 0.2 mL.

### Experimental Design

To determine the effects of host-derived versus environmental microbiome transplants (MT) on cecal microbiome dynamics and pathogen resistance in commercial broiler chicks, we designed a simple factorial experiment (Figure 2A, Table 2) with birds receiving either cecal microbiome transplants (CMT), environmental microbiome transplants (EMT), or PBS control at day-of-hatch. The CMT and EMT inocula were derived and administered as described above and the PBS control was also provided as an oral gavage of 0.2 mL. At 7d post-hatch, half of the birds in each treatment group received a pathogen challenge via oral gavage and the other half remained as controls (Figure 2A). Birds were co-housed until pathogen challenge when they were separated by challenge group. A subset of birds from each treatment group were euthanized and cecal contents removed at the following time points: day-of-hatch, day 7, and day 14 (Figure 1A). For the pathogen challenge, birds in each treatment group were inoculated via oral gavage of 0.2 mL of live *Salmonella enteritis* and *Campylobacter jejuni* cells at an approximate total load of 10^9^ cells for each bacterium. Individual bird weights were recorded as a function of MT type and challenge group (Figure 1B). This experiment was conducted according to the Western University of Health Sciences Institutional Animal care and Use Committee Protocol R15IACUC021.

**Table 2.**
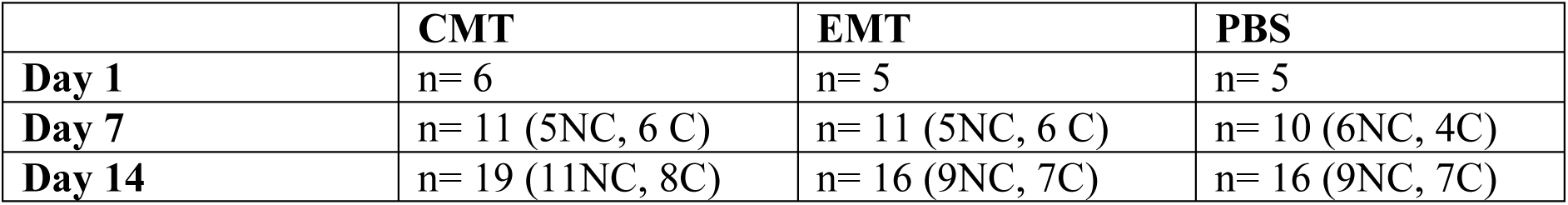
Molecular sequencing replicates. Each replicate represents a cecal community from a euthanized bird. For days 7 and 14, total replicates are subdivided into not challenged (NC) and Challenged (C) groups.

### DNA Extraction and Sequencing

DNA was extracted from ∼100 mg of cecal contents using the MoBio UltraClean Soil DNA extraction kit (Qiagen, Carlsbad, CA) following the manufacture’s protocol. Extracts concentration and quality was checked via spectrophotometry (NanoDrop Products, Wilmington, DE, USA). Amplicons for the V4-V5 hypervariable regions of the 16S rRNA gene were generated via PCR using the 519F (5’-CAG CMG CCG CGG TAA TWC-3’) and 926R (5’-CCG TCA ATT CCT TTR AGG TT-3’) primers following the barcoding scheme of (Faircloth and Glenn, 2012) as detailed elsewhere (Oakley et al., 2013;Oakley and Kogut, 2016). Amplicons were paired-end sequenced on an Illumina MiSeq platform, using a 2×250bp v2 kit, following the manufacturer’s protocol.

### Sequence Analysis

Custom PERL and Unix shell scripts were used to implement portions of the QIIME (Caporaso et al., 2012) and Mothur (Schloss et al., 2009) sequence analyses packages, as described previously (Oakley et al., 2012;Oakley et al., 2013;Oakley and Kogut, 2016). In brief, sequences were trimmed with trimmomatic (Bolger et al., 2014), subsequently merged with Flash (Magoc and Salzberg, 2011), and quality-trimmed (Phred quality threshold of 25) using fastq_quality_trimmer (Blankenberg et al., 2010). Chimera detection was performed with usearch (Edgar et al., 2011) using a type strain database assembled from the SILVA v128 database (Yarza et al., 2010). Taxonomic assignments were performed with usearch against the SILVA database v128 and by the RDP naïve Bayesian classifier against the RDP database (Cole et al., 2014). Sequences were clustered into Operational Taxonomic Units (OTUs) at the RDP genus-level and at 99% sequence similarity with usearch (Edgar et al., 2011).

### Statistical Analyses and Data Summaries

Community analyses were performed in RStudio version 0.98.1091 (Racine, 2012) using the vegan (Oksanen et al., 2015) and phyloseq (McMurdie and Holmes, 2013) R-packages. Briefly, observed community richness was separately assessed for rarefied Genus-level (n= 1012 per sample) and 99% similarity clustered (n=1044 per sample) OTU datasets. Bray-Curtis distances were calculated from the rarefied 99% similarity OTU dataset and used for Principal Coordinate Analyses (PCoA). Differential abundance analyses were performed on abundant taxa (minimum *n* < 100 total reads per OTU) with DESeq2 (Love et al., 2014) using unrarefied experimental subsets, as suggested elsewhere (McMurdie and Holmes, 2014).

## Acknowledgments

Support was provided by the U.S. Poultry & Egg Association, the Western University of Health Sciences College of Veterinary Medicine and the Office of the Vice-President for Research and Biotechnology. The authors thank Mr. George M. Montoya for commenting on a near-final version of this manuscript.

## Author Contributions

BBO, MEB, and NAC conceived the experiment, YD helped with experimental design and project management. ER, JC, and BBO performed the experiment. GAR, JK, and BBO performed bioinformatic analyses. GAR and BBO wrote the manuscript with input from all co-authors.

